# Separation of evolutionary timescales in coevolving species

**DOI:** 10.1101/2023.07.19.549643

**Authors:** Lydia J. Buckingham, Ben Ashby

## Abstract

Many coevolutionary processes, including host-parasite and host-symbiont interactions, involve one species or trait which evolves much faster than the other. Whether or not a coevolutionary trajectory converges depends on the relative rates of evolutionary change in the two species, and so current adaptive dynamics approaches generally either determine convergence stability by considering arbitrary (often comparable) rates of evolutionary change or else rely on necessary or sufficient conditions for convergence stability. We propose a method for determining convergence stability in the case where one species is expected to evolve much faster than the other. This requires a second separation of timescales, which assumes that the faster evolving species will reach its evolutionary equilibrium (if one exists) before a new mutation arises in the more slowly evolving species. This method, which is likely to be a reasonable approximation for many coevolving species, both provides straightforward conditions for convergence stability and is less computationally expensive than traditional analysis of coevolution models, as it reduces the trait space from a two-dimensional plane to a one-dimensional manifold. In this paper, we present the theory underlying this new separation of timescales and provide examples of how it could be used to determine coevolutionary outcomes from models.

## Introduction

Coevolution is ubiquitous in nature and has been implicated in many biological phenomena, including diversification (Futuyma and Agrawal, 2009; Paterson et al., 2010; Thompson, 2009) and the evolution of sex (Hamilton et al., 1990; Morran et al., 2011). Coevolving species may be mutualistic, such as host-symbiont (Janzen, 1966; Limborg and Heeb, 2018) or plant-pollinator (Johnson and Anderson, 2010) relationships; antagonistic, such as host-parasite (Flor, 1956; Marshall and Fenner, 1960, 1958) (including brood parasitism (Feeney et al., 2014)) or predator-prey (Heiling and Herberstein, 2004; Vermeij and Covich, 1978) (including herbivory (Ehrlich and Raven, 1964)) relationships; or competitive (Leger and Espeland, 2010). Coevolution can lead to rapid reciprocal adaptations, but in many cases species evolve at very different rates, especially when interactions occur across trophic levels (e.g. plant or animal interactions with microbial species (Drew et al., 2021)).

For example, bacterial or fungal symbionts typically evolve much faster than their hosts (Moran et al., 1995) and viral or bacterial parasites have generation times and population sizes that are orders of magnitude faster or larger than their plant or animal hosts, setting the stage for much faster adaptive evolution (Bliven and Maurelli, 2016; Elena et al., 2008). This includes RNA viruses of humans, such as coronaviruses and influenza viruses, which typically have generation times measured in hours with tens of billions of virions per infection, whereas humans have generation times measured in decades with fewer individuals in the entire human population than virions in a single infected host (Bar-On et al., 2020; Sender et al., 2021). Of course, vertebrates can often keep pace with parasites through adaptive immunity, which in many cases is the more pertinent driver of parasite evolution over shorter timescales, but over longer timescales parasites can drive evolution in vertebrate populations, leading to coevolution between a host and a much faster evolving microbe. Beyond host-microbe interactions, contrasting generation times and population sizes occur in a wide range of other ecological relationships, including plant-herbivore (e.g. insect herbivores have much shorter lifespans and so evolve much faster than long-lived trees (Edmunds and Alstad, 1978; Karban, 1989)), predator-prey (e.g. krill and whales (Jarman, 2001; Meredith et al., 2013; Pyenson, 2017)) and competitive interactions (e.g. fast evolving algae compete for space with more slowly evolving corals (Swierts and Vermeij, 2016)).

One can employ a variety of methods to model coevolutionary dynamics theoretically (Ashby et al., 2019; Buckingham and Ashby, 2022), including population genetics (Day and Gandon, 2007), quantitative genetics (Nuismer et al., 2007), adaptive dynamics (Dieckmann and Law, 1996; Geritz et al., 1998; Kisdi, 2006; Metz et al., 1996) and oligomorphic dynamics (Sasaki and Dieckmann, 2011). Adaptive dynamics (also known as evolutionary invasion analysis) is especially useful for modelling the long-term evolutionary dynamics of quantitative traits, as it naturally incorporates population (ecological) dynamics and is relatively straightforward to implement. A key assumption of this method is a separation of ecological and evolutionary timescales, so that the system reaches its ecological attractor (usually a stable equilibrium or limit cycle (Best and Ashby, 2023)) before a new mutation arises. This greatly simplifies the analysis because ecological dynamics (for instance, changes in population density) can be considered separately to evolutionary (trait) dynamics. However, important feedbacks remain between the ecological and evolutionary dynamics that play a fundamental role in shaping coevolution (Ashby et al., 2019). Separating evolutionary and ecological timescales is also a reasonable approximation for many real biological systems, as evolutionary dynamics are often, but not always (Pelletier et al., 2009), much slower than ecological dynamics.

Here, we generalise and elaborate on a recent method for separating evolutionary timescales in coevolving species (Ashby and Farine, 2022). We propose that, for systems where one species typically evolves much faster than the other (e.g., due to stark differences in generation times, population sizes or mutation rates), one can reasonably introduce a second separation of timescales between the evolutionary dynamics of the two species. This greatly simplifies the analysis by collapsing coevolutionary dynamics in a two-dimensional plane into a one-dimensional manifold. In this paper, we will describe this method for modelling the coevolution of two species and consider how the results compare to existing adaptive dynamics methods.

## Methods

We begin by outlining the classical adaptive dynamics framework for two coevolving species, before introducing an approximation using a separation of evolutionary timescales between species. For convenience, we will use the example of a host-parasite system throughout, although these methods may be applied to many other coevolving species. Thus, “parasite” may be used interchangeably with “fast evolving species” and “host” with “slowly evolving species”. We will only consider the case of two species, each with a single evolving trait, but this method could also be applied to two coevolving traits within the same species, as long as one evolves much faster than the other, or to systems with more than two coevolving traits. Note that the rates of evolutionary change in each species will depend on the size and frequency of mutations, population sizes and the selection gradient. Here, when we refer to a fast evolving species (e.g., a parasite), we mean that these factors combine to give a much higher rate of evolutionary change than in the slowly evolving species.

### Adaptive dynamics for coevolving species

The adaptive dynamics framework for two coevolving, asexual species has been described in detail elsewhere (Dieckmann and Law, 1996; Geritz et al., 1998; Kisdi, 2006; Metz et al., 1996), so here we shall only give a brief overview. The framework makes two crucial assumptions. First, that there is a separation of ecological and evolutionary timescales, such that a mutation may only occur after the ecological dynamics have reached an attractor (e.g. an equilibrium or a limit cycle). In other words, selection acts relatively quickly so that the fate of a mutant is determined before another mutant arises. Second, mutations are assumed to have small additive effects, which means that traits are modelled as continuous with mutants phenotypically similar to their progenitors. Note that even though traits are modelled as continuous, mutation effect sizes are assumed to be small but not infinitesimal. This allows for outcomes such as evolutionary branching, which cannot occur if mutations have infinitesimal effects (Geritz et al., 1998).

For (initially) monomorphic host (H)and parasite (*p)*populations, with evolvable traits *h* and *p* respectively, the invasion fitness of a rare mutant *h*_*m*_ or *p*_*m*_ is given by a function of the form *w*_*H*_(*h*_*m*_, *h, p*)or *w*_*p*_*(p*_*m*_, *h, p*), where the current resident population determines the “environment” that the mutant experiences. The traits evolve in the direction of their respective fitness gradients, 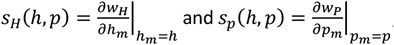, until a pair of *co-singular strategies* (*h*^*^, *p*^*^) are reached, which satisfy *s*_*H*_(*h*^*^, *p*^*^)=*s*_*p*_*(h*^*^, *p*^*^)=0. The stability of a pair of co-singular strategies is determined both in terms of *evolutionary stability*, which tells us whether a rare mutant can invade, and by *convergence stability*, which tells us whether trait values that are sufficiently close to (*h*^*^, *p*^*^)will tend towards this point in the long-term. A pair of co-singular strategies is evolutionarily stable for the host if:

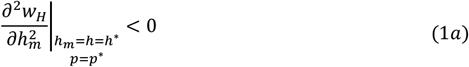

and is evolutionarily stable for the parasite if:

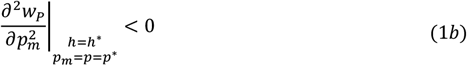

Though convergence stability is relatively straightforward to calculate when there is only one evolving trait, it is more difficult when there are two or more traits because convergence stability depends on the relative rates of evolutionary change (Geritz et al., 1998; Kisdi, 2006). In simple systems, one may derive sufficient, but not necessary, conditions for convergence. For example, *strong convergence stability* (Leimar, 2009) is the property whereby every coevolutionary trajectory with sufficiently small mutational steps will converge to a pair of co-singular strategies. Specifically, co-singular strategies are strongly convergence stable if the following three conditions all hold:

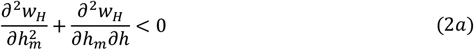

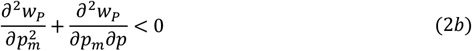

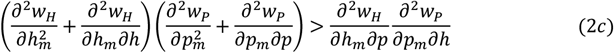

where all derivatives are evaluated at the co-singular strategies (*h*_*m*_ =*h* =*h*^*^ and *p*_*m*_ =*p* =*p*^*^). Since this is a sufficient, but not necessary, set of conditions for convergence to (*h*^*^, *p*^*^), a pair of co-singular strategies may not be strongly convergence stable even if many coevolutionary trajectories still converge to it. A pair of co-singular strategies which is convergence stable and evolutionarily stable for both the host and parasite is called a *co-continuously stable strategy* (co-CSS).

### Separation of host and parasite evolutionary timescales

In systems where the rate of evolutionary change in one species (e.g., a parasite) is much faster than the other (e.g., a host), we can make a second separation of timescales, between host and parasite evolutionary dynamics. We therefore have ecological dynamics occurring on a much faster timescale than parasite evolution, which in turn occurs on a much faster timescale than host evolution. This is a reasonable assumption for many host-parasite interactions, where parasite population sizes and generation times may be many orders of magnitude larger and shorter, respectively, than in the host. We therefore assume that, for a given value of the host trait *h*, the parasite reaches an evolutionary attractor (if one exists) before the next host mutation occurs.

A singular strategy for the parasite (if one or more singular strategies exist) as a function of the host trait 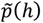, is now found by solving:

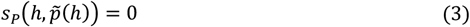

Such a singular strategy is then evolutionarily stable for the parasite if:

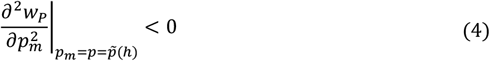

and is convergence stable if:

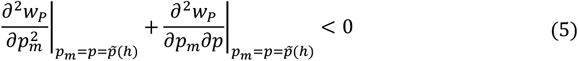

Note that these conditions depend on the host trait value, *h*, but do not depend on the rate of evolutionary change in the host, due to the second separation of timescales.

For a given value of the host trait, the long-term trait dynamics of a fast-evolving parasite may be: (1) a monomorphic population that tends to a continuously stable strategy or a maximum or minimum trait value; (2) a polymorphic population arising from evolutionary branching into two or more traits, which tend to a set of continuously stable strategies, and/or maximum or minimum trait values; (3) directional selection in one or more trait values that tend to positive or negative infinity; (4) fluctuating selection in one or more trait values (e.g., due to seasonality in the ecological dynamics); (5) drift or (6) extinction of one or both populations. Here, we focus on scenarios 1-2, as scenarios 3-5 represent non-equilibrium evolutionary dynamics for the parasite (and may therefore require consideration of Floquet exponents (Ferriere and Fox, 1995; Ferriere and Gatto, 1995; Geritz et al., 2007; Klausmeier, 2008) or stochasticity) and scenario 6 is trivial as extinction of one or both species prevents further coevolution.

### SCENARIO 1: MONOMORPHIC PARASITE POPULATION

Suppose that the parasite remains monomorphic and tends to either a continuously stable strategy 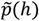 is evolutionarily stable and convergence stable), or a maximum or minimum trait value (if the fitness gradient at a boundary points towards the boundary). We denote the evolved parasite trait by 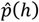 (note that in the case of a CSS,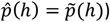).

The invasion fitness of a rare host mutant is then given by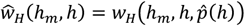, with fitness gradient 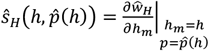 The host singular strategy therefore satisfies 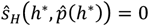 =0, and is evolutionarily stable if:

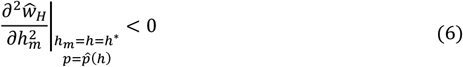

and is convergence stable if:

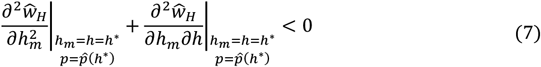

Equation (7) can be re-written in terms of the original host invasion fitness, *w*_H_, as follows (for derivation see *Supplementary Materials*):

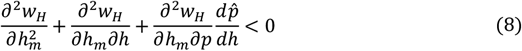

where all terms are evaluated at *h*_*m*_ =*h* =*h*^*^ and *p* =*p*^*^.

Therefore, a co-singular strategy is (at least locally) convergence stable when the parasite evolves much faster than the host if both conditions (5) and (8) hold at the co-singular strategy. These two conditions can be formulated in a number of ways, and in fact are equivalent to conditions (2b) and (2c), which are two of the conditions required for strong convergence stability (see *Supplementary Materials* equations (S6) to (S13) for derivation). That is, all three of the conditions (2a) to (2c) must hold for a co-singular strategy to be strong convergence stable (for all rates of evolution), whereas only conditions (2b) and (2c) need hold for the co-singular strategy to be convergence stable when the parasite evolves much faster than the host. These findings concur with the results of Leimar (2009), whose conditions for convergence stability can also be shown to reduce to our conditions (2b) and (2c) in the case where the parasite evolves arbitrarily quickly relative to the host (see *Supplementary Materials*).

This formulation of our conditions makes it clear that, if (*h*^*^, *p*^*^)is a co-CSS (or is a boundary point that behaves like a co-CSS) in the classical adaptive dynamics framework, then it is also a co-CSS using our separation of host and parasite evolutionary timescales method. This makes sense because strong convergence stability implies that all coevolutionary trajectories close to (*h*^*^, *p*^*^)converge to the co-singular point, including those where the parasite evolves much faster than the host. The reverse implication does not always hold, however; just because coevolutionary trajectories where the parasite has a much faster rate of evolutionary change than the host converge to a particular point, it does not mean that all other trajectories will.

Condition (8) may be interpreted geometrically in terms of a phase plane (see Fig. 1). The sum of the first two terms, 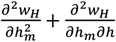, represents the rate of change of the host component of the fitness gradient arrows as we move horizontally through the co-singular strategy. The Term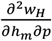 represents the rate of change of the host component of the arrows as we move vertically through the co-singular strategy. The term 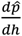 represents the gradient of the parasite nullcline at the co-singular strategy. Therefore, if the fitness gradient arrows point to the right on the left of the co-singular strategy and point to the left on the right of the co-singular strategy then a negative slope of the parasite nullcline promotes convergence stability (and vice versa).

**Fig. 1:**
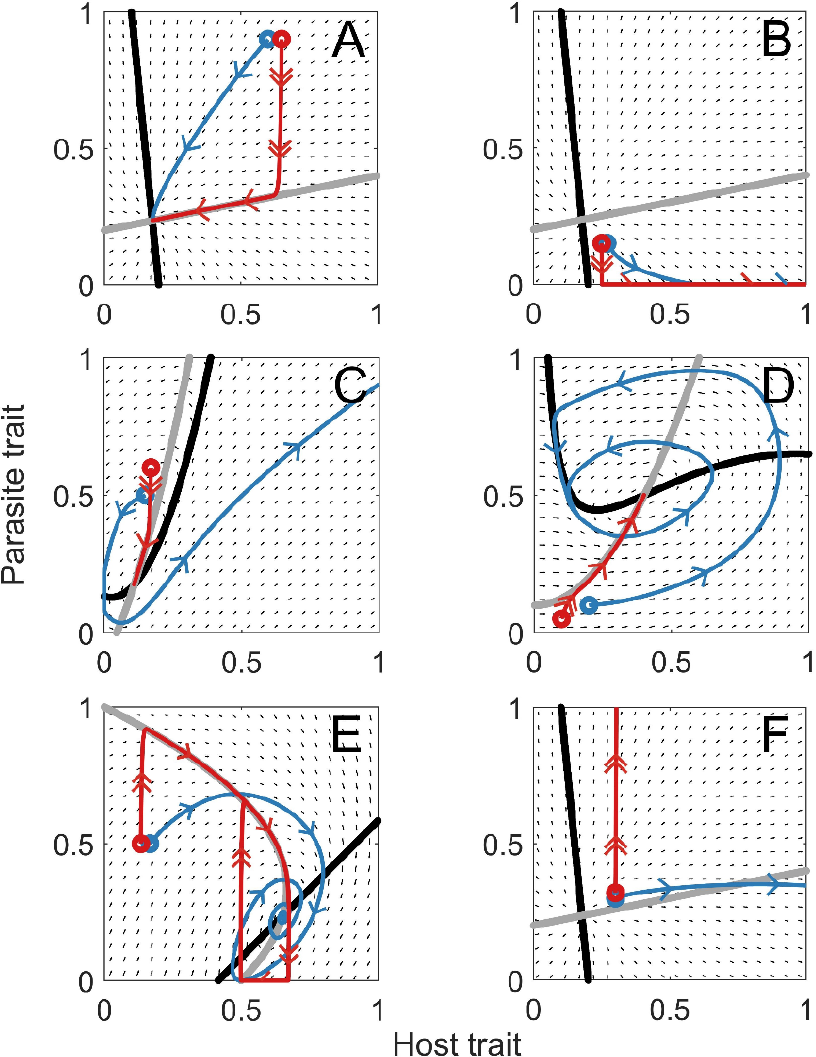
Phase planes showing coevolutionary trajectories when the two species have comparable rates of evolutionary change (blue) and when the parasite evolves much faster than the host (red). Double arrows indicate periods of rapid evolution and single arrows indicate slower rates of evolution. In (A) and (B), the long-term outcome does not depend on the relative rates of evolutionary change, with both trajectories either converging to (A) or diverging from (B) the co-singular strategy. In (C) and (D), the traits converge to the co-singular strategy when the parasite evolves much faster than the host, but when the two species have comparable rates of evolutionary change, the traits may increase indefinitely (C) or cycle (D). In (E), both traits converge when the species evolve at comparable rates but cycle when the parasite evolves much faster than the host. In (F), both trajectories evolve away from the co-singular strategy, but towards different evolutionary endpoints. The host nullcline is shown in black and the parasite nullcline in grey. Circles mark the initial values of the evolving traits for the given trajectories. Note that these phase planes were generated manually to demonstrate a range of possible outcomes.

To illustrate the effects of separating evolutionary timescales, consider the phase planes in Fig. 1, where the co-singular point is found at the intersection of the nullclines for the host (black) and parasite (grey) fitness gradients. If the parasite evolves much faster than the host (red), then the coevolutionary trajectory rapidly moves to (or directly away from) the parasite nullcline and then follows the nullcline (or an edge of the phase plane). Convergence stability is determined by whether the trajectories move towards (Fig. 1A) or away from (Fig. 1B) the parasite nullcline, and then (if they do converge to the nullcline) whether they move along the parasite nullcline towards or away from the co-singular strategy.

However, even if all trajectories converge to the co-singular strategy when the parasite evolves much faster than the host, all trajectories may evolve away from the co-singular strategy (or cycle and therefore never approach it) when the rates of evolutionary change in both species are comparable (Fig. 1C,D; see also *Example 1* below). It is also possible for a co-singular strategy to be an attractor of any mutational path with comparable rates of evolutionary change but not of any mutational path where the parasite evolves much faster than the host (such a co-singular strategy is not strong convergence stable, because it is not an attractor of all mutational paths). This can lead to a case where the parasite trait is initially convergence stable, but as the host trait subsequently evolves the trajectory moves to a region where the parasite is not convergence stable. This can generate stable cycles in cases where cycling is not seen for comparable rates of evolutionary change (Fig. 1E). A co-singular strategy may also be convergence unstable for all mutational paths, but the relative rates of evolutionary change may determine, for instance, whether both traits increase indefinitely or whether one increases while the other falls (Fig. 1F).

What can we say in general about the nature of the phase plane for systems with disparate and similar rates of evolutionary change, and the resulting coevolutionary trajectories? Intuitively, when the parasite nullcline is an attractor and has a relatively shallow gradient near a co-singular point, a relatively small change in the host phenotype does not lead to a sudden, large change in the parasite phenotype. Coevolutionary trajectories therefore gradually move along the parasite nullcline when the latter species evolves much faster, converging to the same co-CSS as in classical adaptive dynamics (Fig. 1A). In contrast, when the parasite nullcline has a relatively steep gradient near a co-singular point, a relatively small change in the host phenotype leads to a sudden large shift in the parasite phenotype, potentially leading to a difference in the long-term dynamics (Fig. 1C-E). Examining the gradient and convergence stability of the parasite nullcline will therefore give a strong indication as to whether the long-term dynamics of the system with disparate rates of evolutionary change generalise to when rates of evolutionary change are more comparable.

### SCENARIO 2: POLYMORPHIC PARASITE POPULATION

If the singular strategy 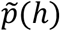 is convergence stable (equation 5) but not evolutionarily stable (equation 4), then the parasite will branch into two traits (note that since we are essentially dealing with single trait evolution due to the separation of host and parasite timescales, mutual invasibility is implied (Geritz et al., 1998)). As a branching point is not an evolutionary endpoint, we must determine what eventually happens to the two parasite traits before we can consider the fate of a host mutation. We must therefore form a three species system with one host and two parasite species and modify the parasite invasion fitness to account for the two resident phenotypes, *p*_1_ and 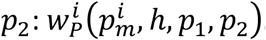 . At this point, the analysis proceeds as in the monomorphic scenario to determine if there is a pair of co-singular strategies for the parasite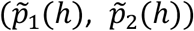, given the current host trait value. If this point exists and is evolutionarily stable, then the invasion fitness of a rare host mutant is given by 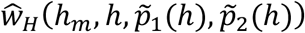 and the analysis follows the same logic as above. If the parasite branches again, then one must modify the parasite invasion fitness to account for three resident phenotypes, and so on.

## Examples

### Example 1: Differences in convergence stability

We consider the model of host-parasite coevolution proposed by Best *et al*. (2010). Susceptible hosts (with density *S*) reproduce at an underlying rate, *a*, subject to density-dependent competition, *q* (infected hosts do not reproduce). Disease transmission is also density-dependent with transmission rate *β*. Hosts die naturally at a constant rate, *b*, but infected hosts (with density *I*) also experience disease-induced mortality, *α*. We assume that there is no recovery from infection for simplicity. The ecological dynamics are modelled using the following system:

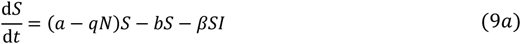

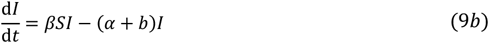

We assume that the hosts and parasites have evolvable traits *h* and *p* respectively, which influence their life-histories through the trade-off functions *a* =*a*(*h*)and *β* =*β*(*h, p*)(as in (Best et al., 2010)). We consider the following functional forms for the trade-offs (for simplicity of calculations, these are different functional forms to the trade-offs used by Best *et al*. (2010) but have similar shapes and properties):

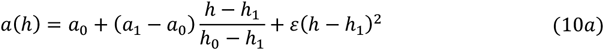

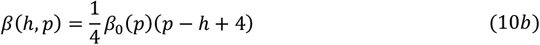

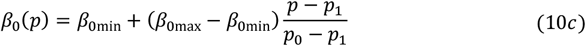

We assume that *h*_0_ ≤ *h* ≤ *h*_1_ and *p*_0_ ≤ *p* ≤ *p*_1_ (where *h*_0_ =*p*_0_ =0 and *h*_1_, *p*_1_ ≤ 4 to ensure that *β* ≥ 0). We can see that the transmission rate of the parasite depends on the difference between the host and parasite traits (*p − h*) and so these traits can be interpreted as host resistance to infection (which comes at a cost to host reproduction, *a*) and parasite infectivity (which increases baseline transmissibility, *β*_0_). We let *Φ* represent the relative mutation rate of the parasite trait (the parasite mutates *Φ* times faster than the host).

Analytical calculations (given in the *Supplementary Materials*) show that, at any co-singular strategy, the parasite is always evolutionarily stable, whereas the host may be evolutionarily stable or unstable, depending on parameter values (hence branching may occur in the host trait). The parasite is always convergence stable when evolving quickly relative to the host, no matter the value of the host trait, and the host may also be convergence stable (depending on parameter values). Strong convergence stability of the co-singular strategy is not guaranteed.

These conditions mean that we can find parameter values (see Table S1) for which the co-singular strategy is not strong convergence stable (and moreover the traits do not converge when the rates of evolutionary change in the host and parasite are comparable), but where both host and parasite traits are convergence stable when the parasite is evolving quickly relative to the host (Fig. 2). In this particular case, the host and parasite traits converge to a branching point when the parasite evolves much faster than the host (Fig. 2A,B). When the host and parasite have comparable rates of evolutionary change, however, their traits do not converge to their co-singular strategy and instead both tend to zero (Fig. 2C,D).

**Fig. 2:**
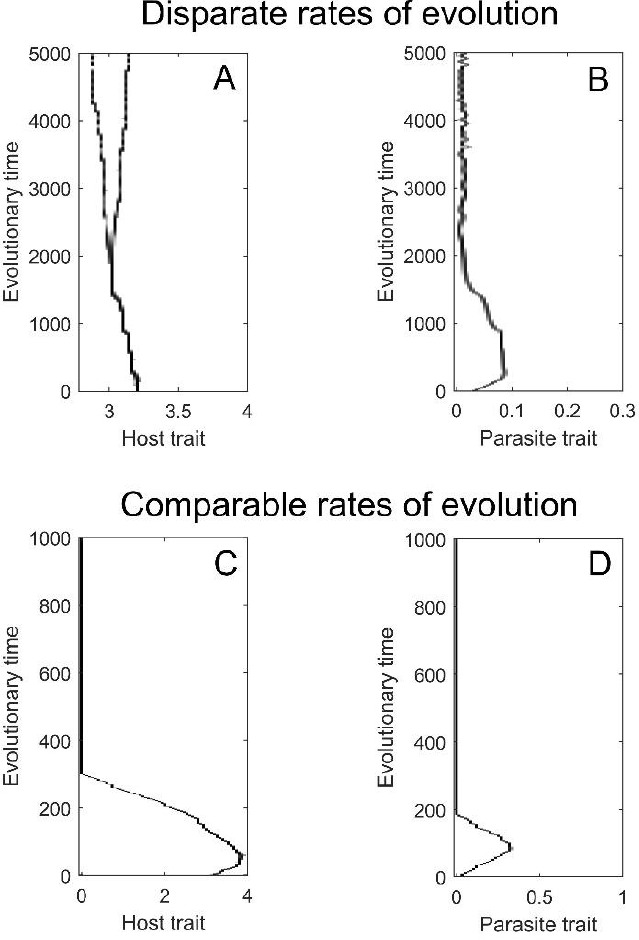
Simulations showing the case where trajectories converge to the co-singular strategy when the parasite evolves much faster than the host (A & B; *Φ* =*1*00) but where the trajectories do not converge to the co-singular strategy when the host and parasite have comparable rates of evolutionary change (and hence the co-singular strategy is not strong convergence stable; C & D; *Φ* =*1*). Parameters used are as in Table S1.

This example demonstrates that considering only the case of comparable rates of evolutionary change, or considering sufficient (but not necessary) conditions like strong convergence stability, may miss key predictions. In this case, both methods would conclude that the co-singular strategy is not convergence stable, even though coevolutionary trajectories do converge whenever the parasite evolves sufficiently faster than the host. Therefore, if there is reason to believe that the parasite should evolve much faster than the host, separating the evolutionary timescales of the two species provides a more realistic prediction.

### Example 2: Differences in branching

Svennungsen & Kisdi (2009) consider the evolution of parasite virulence (disease-induced mortality, *α*) subject to a trade-off with transmissibility, *β*(*α*)(Svennungsen and Kisdi, 2009). Their model assumes that all hosts reproduce at a constant rate, *b*, and experience both density-independent and density-dependent intrinsic mortality, given by the parameters *A* and *B*, respectively. Disease transmission is density-dependent and infected hosts experience disease-induced mortality. These ecological dynamics are described by the following system:

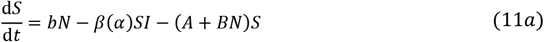

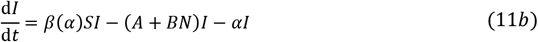

where *N* is the total host population density and the trade-off function is given by:

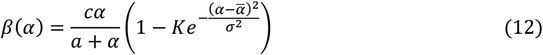

This model has been shown to exhibit branching in virulence for a variety of parameter values (Svennungsen and Kisdi, 2009).

We can extend this model to consider coevolution with host resistance, *r*, subject to a trade-off with reproduction, *b*. To do this, we introduce the following functions:

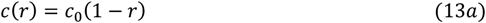

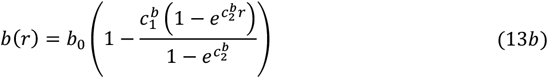

with *c* =*c*(*r*)in equation (12) corresponding to the level of host susceptibility and *b* =*b*(*r*)in equation (11a) corresponding to the reproduction cost associated with resistance. We let *Φ* represent the relative mutation rate of the parasite trait (the parasite mutates *Φ* times faster than the host).

The resistance trait, *r*, varies between zero (no resistance) and one (full resistance). When *r* =0, we regain the original system with *b* =*b*_0_ and *c* =*c*_0_. As host resistance increases, the transmission rate between hosts, *β*, falls until complete resistance is reached (when *r* =*1*, no transmission occurs). As resistance increases, host reproduction also falls (reaching a minimum level *b* =*b*_0_(*1 − c*^b^)when *r* =*1*). A summary of model parameters and variables is given in Table S2.

If host resistance is initially low enough and evolves sufficiently slowly relative to the parasite, then we know that parasite virulence can branch into two distinct phenotypes (Svennungsen and Kisdi, 2009). However, if the host and parasite have comparable rates of evolutionary change then branching does not necessarily occur for the same parameter values (Fig. 3). Even though the conditions for evolutionary stability of a co-singular strategy are the same for both the traditional adaptive dynamics framework (equation 1) and our new separation of evolutionary timescales method (equations 4 and 6), the incidence of branching does not need to be the same for all rates of evolutionary change. This is because it is possible for the only co-singular strategy to be a co-CSS (and hence when rates evolutionary change are comparable both species will evolve towards it and not branch) but for some points along the parasite nullcline far from the co-CSS to be evolutionarily unstable (and hence when the parasite evolves sufficiently fast, it may branch before the host has mutated). Whether the rates of evolution are disparate or comparable can also have a significant quantitative effect on the evolutionary equilibrium of the traits (e.g. Fig. 3A,C).

**Fig. 3:**
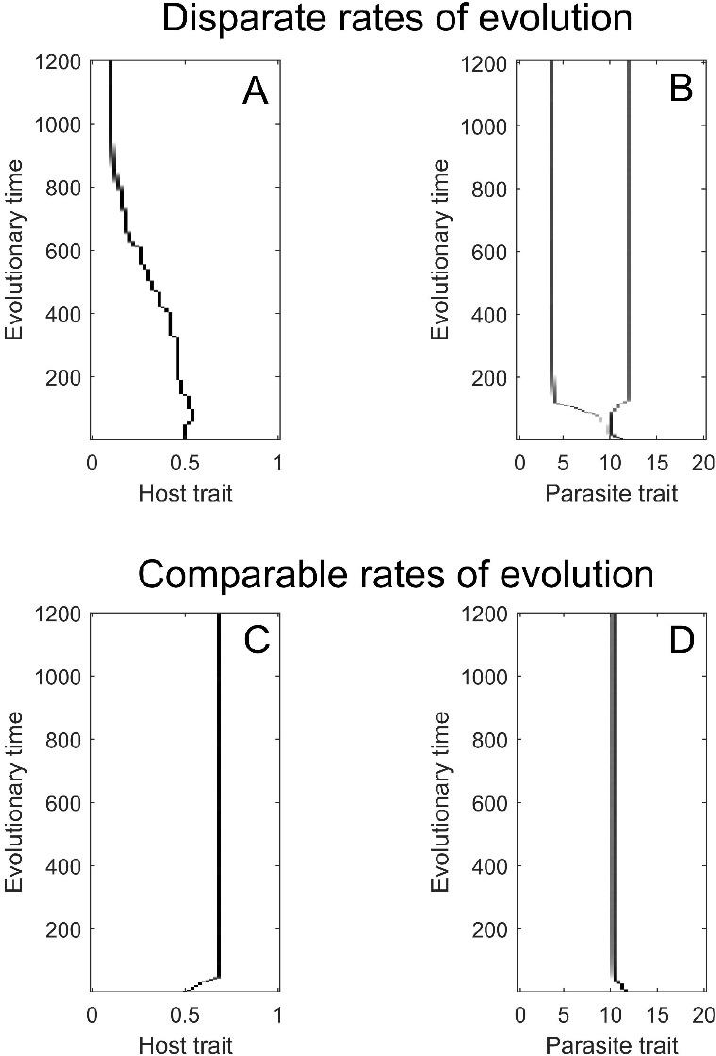
Simulations showing the case where branching occurs when the parasite evolves much faster than the host (A & B; *Φ* =20) but where the co-singular strategy is continuously stable (a co-CSS) when the host and parasite have comparable rates of evolutionary change (C & D; Φ =*1*). Parameters used are as in Table S2.

## Discussion

In many coevolving systems, especially those that involve more than one trophic level, evolution may proceed at very different rates within each species (Moran et al., 1995). This has important implications from a theoretical perspective, because coevolutionary convergence stability in adaptive dynamics depends on the relative rates of evolutionary change in the coevolving species (Dieckmann and Law, 1996; Marrow et al., 1996; Matessi and Pasquale, 1996). In this paper, we have shown how modelling the coevolution of two species on separate evolutionary timescales can greatly simplify model analysis and provide results which are more closely tailored to systems where species have contrasting rates of evolutionary change.

The relative rates of evolutionary change of two coevolving species can have a significant impact on coevolutionary outcomes (Dieckmann and Law, 1996; Marrow et al., 1996; Matessi and Pasquale, 1996). While some studies do consider the effect of varying relative rates of evolutionary change on coevolutionary dynamics (Best et al., 2010), many do not. Instead, models that use the adaptive dynamics framework typically either assume arbitrary rates of evolutionary change (e.g., (Best et al., 2008; Law et al., 2001; Rafaluk-Mohr et al., 2018)) or restrict analysis to cases where there is strong convergence stability (i.e., convergence stability holds for all rates of evolutionary change) (Best et al., 2009). Comparable rates of evolutionary change may be a good approximation for some biological systems (Nair et al., 2019; Naureen et al., 2020; Pollock et al., 2021), but in cases where species have very different evolutionary timescales (as is often true in host-microbe relationships), if similar rates are used in modelling then theoretical results may represent a significant departure from the real dynamics of natural populations.

Our approach considers the limit of rapid evolution in one species, where it reaches an evolutionary endpoint (if one exists) before a mutation arises in the other species. This additional separation of timescales is an extension to the standard separation of ecological and evolutionary timescales in adaptive dynamics, but also parallels other examples of separate timescales. For example, Fortelius *et al*. considered the effects of slow climatic change on evolution by introducing an additional separation of evolutionary and geological timescales into an adaptive dynamics model (Fortelius et al., 2015a, 2015b). Our approach, however, is the first to explicitly consider separating two evolutionary timescales based on different rates of adaptation across species. This method had previously been applied to a model of the coevolution of animal sociality and parasite virulence (Ashby and Farine, 2022) but here we have generalised the method and provided greater analytical details (e.g., conditions for convergence stability). Naturally, this method will provide a better approximation to biological systems which have greater differences between the rates of evolutionary change of the two species. Furthermore, the approximation is likely to generalise to comparable rates of evolutionary change when the nullcline for the fast evolving species has a relatively shallow gradient and is convergence stable.

If rates of evolutionary change are unknown for a given system, then one can only derive necessary or sufficient conditions for convergence stability or branching (Kisdi, 2006; Leimar, 2009). For instance, strong convergence stability is useful for proving that all coevolutionary trajectories tend to a particular point in trait space, but many of these trajectories will typically be unrealistic (e.g., parasites evolving much slower than their hosts). It is generally not important to know whether a particular result holds for all rates of evolutionary change, only whether it will hold for the typical rates of evolutionary change of the species in question (or the most likely rates of evolutionary change if they are unknown). In cases where one species is likely to evolve much faster than the other (e.g., host-parasite systems), separating their evolutionary timescales not only simplifies the analysis but may provide more realistic results by approximating the rapid evolution of one species.

There are two key advantages of our separation of evolutionary timescales method. First, our approximation gives a more straightforward definition of convergence stability when one species is known to evolve much faster than the other, rather than one which relies on relative rates of evolutionary change. Second, it is less computationally expensive because we are effectively only considering a one-dimensional manifold within a two-dimensional trait space. Rather than having to calculate the fitness gradients for every possible pair of traits, we only need to find them along the nullcline of the faster-evolving species. Together, these advantages mean that our method greatly simplifies and speeds up the analysis of coevolutionary models. Even if one is generally interested in a wide range of rates of evolutionary change, our method offers an efficient means of testing what happens when differences in these rates are large. Furthermore, the methods proposed herein could be readily extended to consider additional separations of timescales across trophic levels (e.g., hosts, parasites and hyperparasites (Wood and Ashby, 2023)) or the coevolution of multiple traits in each species. For instance, multiple traits in a host organism (such as different resistance or tolerance mechanisms) may evolve on similar timescales while multiple traits in the parasite (such as mortality and sterility virulence) evolve much more quickly. In this case we could make the assumption that all parasite traits reach their co-evolutionary endpoints before any new mutations arise in the host.

However, this method is clearly not appropriate for all coevolving species. We have shown that, in some cases, separating evolutionary timescales produces qualitatively and quantitatively different evolutionary outcomes to when species have comparable rates of evolutionary change. As such, this method is more suitable than others for modelling systems where one species evolves much faster than the other, but is not suitable for modelling systems in which the species evolve on similar timescales (e.g., in microbial communities, such as bacteria-phage coevolution). Still, our method offers a relatively straightforward test for determining whether results which assume comparable (or arbitrary) rates of evolutionary change still apply when one species evolves much faster than the other, and so will be especially beneficial in scenarios restricted to plants/animals coevolving with microorganisms (e.g., parasite-mediated sexual selection/conflict (Ashby, 2020; Ashby and Boots, 2015; Hamilton and Zuk, 1982; Pirrie et al., 2022; Wardlaw and Agrawal, 2019)).

Overall, it is important to consider relative rates of evolutionary change when modelling the coevolution of two species. If rates of evolutionary change are likely to be similar, then it is important that the analysis is not over-simplified and the approximation presented herein may not be appropriate. However, if one species is known (or is likely) to evolve much faster than the other, then our method can greatly simplify the analysis and ensure more realistic predictions.

## Data availability

Source code is available in the *Supplementary Materials* and at: https://github.com/ecoevotheory/Buckingham_and_Ashby_2023c.

## Acknowledgements

We thank Alex Best for helpful comments on the manuscript. Ben Ashby is supported by the Natural Environment Research Council (grant no. NE/V003909/1). This research was generously supported by a Milner Scholarship PhD grant to Lydia Buckingham from The Evolution Education Trust. We acknowledge the support of the Natural Sciences and Engineering Research Council of Canada (NSERC). Nous remercions le Conseil de recherches en sciences naturelles et en génie du Canada (CRSNG) de son soutien. PIPPS receives funding from the BC Ministry of Health.

