## Supplementary Materials for "Separation of evolutionary timescales in coevolving species"

### Separation of Evolutionary Timescales in Coevolving Species: Supplementary Materials

#### HOST CONVERGENCE STABILITY FOR SEPARATE EVOLUTIONARY TIMESCALES

By the standard condition for convergence stability of a single evolving trait, we know that the host trait is convergence stable if and only if:

$$\left.\frac{\partial^2 \hat{w}_H}{\partial h_m^2}\right|_{\substack{h_m=h=h^* \\ p=\hat{p}(h^*)}} + \left.\frac{\partial^2 \hat{w}_H}{\partial h_m \partial h}\right|_{\substack{h_m=h=h^* \\ p=\hat{p}(h^*)}} < 0 \qquad (S1)$$

Now, $\hat{w}_H(h_m, h) = w_H(h_m, h, \hat{p}(h))$ and so, by the chain rule:

$$\frac{\partial \hat{w}_H}{\partial h} \equiv \frac{\partial w_H}{\partial h} + \left.\frac{\partial w_H}{\partial p}\right|_{p=\hat{p}(h)} \frac{d\hat{p}}{dh} \qquad (S2)$$

Differentiating again with respect to the mutant host trait then gives:

$$\frac{\partial^2 \hat{w}_H}{\partial h_m \partial h} \equiv \frac{\partial^2 w_H}{\partial h_m \partial h} + \left.\frac{\partial w_H}{\partial h_m \partial p}\right|_{p=\hat{p}(h)} \frac{d\hat{p}}{dh} \qquad (S3)$$

since $\hat{p}$ is not a function of $h_m$. We also know that:

$$\frac{\partial^2 \hat{w}_H}{\partial h_m^2} = \frac{\partial^2 w_H}{\partial h_m^2} \qquad (S4)$$

Evaluating expressions (S3) and (S4) at $h_m = h = h^*$ and $p = p^*$ and substituting back into equation (S1) then tells us that equation (S1) is equivalent to:

$$\frac{\partial^2 w_H}{\partial h_m^2} + \frac{\partial^2 w_H}{\partial h_m \partial h} + \frac{\partial^2 w_H}{\partial h_m \partial p}\frac{d\hat{p}}{dh} < 0 \qquad (S5)$$

evaluated at $h_m = h = h^*$ and $p = p^*$.

#### FORMULATIONS OF CONVERGENCE CONDITIONS FOR SEPARATE EVOLUTIONARY TIMESCALES

At a co-singular strategy, we know that:

$$s_P(h, p) = 0 \Leftrightarrow p = \tilde{p}(h) \qquad (S6)$$

We can differentiate this equation implicitly with respect to $h$ and evaluate at $p = \tilde{p}(h)$ to give:

$$\frac{\partial s_P}{\partial h} + \frac{\partial s_P}{\partial p}\frac{d\tilde{p}}{dh} = 0 \qquad (S7)$$

We know that $s_p(h,p) = \left.\frac{\partial w_P}{\partial p_m}\right|_{p_m=p}$ and so this can be re-written as:

$$\frac{\partial^2 w_P}{\partial p_m \partial h} + \left(\frac{\partial^2 w_P}{\partial p_m \partial p} + \frac{\partial^2 w_P}{\partial p_m^2}\right)\frac{d\tilde{p}}{dh} = 0 \qquad (S8)$$

where everything is evaluated at $p_m = p = \tilde{p}(h)$. We can then re-arrange this equation to:

$$\frac{\partial^2 w_P}{\partial p_m \partial h} = -\left(\frac{\partial^2 w_P}{\partial p_m \partial p} + \frac{\partial^2 w_P}{\partial p_m^2}\right)\frac{d\tilde{p}}{dh} \qquad (S9)$$

This condition always holds at a co-singular strategy. Now assume that we have a co-singular strategy which satisfies conditions (2b) and (2c) from the main text. That is:

$$\frac{\partial^2 w_P}{\partial p_m^2} + \frac{\partial^2 w_P}{\partial p_m \partial p} < 0 \qquad (2b)$$

$$\left(\frac{\partial^2 w_H}{\partial h_m^2} + \frac{\partial^2 w_H}{\partial h_m \partial h}\right)\left(\frac{\partial^2 w_P}{\partial p_m^2} + \frac{\partial^2 w_P}{\partial p_m \partial p}\right) > \frac{\partial^2 w_H}{\partial h_m \partial p}\frac{\partial^2 w_P}{\partial p_m \partial h} \qquad (2c)$$

Then we can substitute equation (S9) into (2c) to get:

$$\left(\frac{\partial^2 w_H}{\partial h_m^2} + \frac{\partial^2 w_H}{\partial h_m \partial h}\right)\left(\frac{\partial^2 w_P}{\partial p_m^2} + \frac{\partial^2 w_P}{\partial p_m \partial p}\right) > -\frac{\partial^2 w_H}{\partial h_m \partial p}\left(\frac{\partial^2 w_P}{\partial p_m \partial p} + \frac{\partial^2 w_P}{\partial p_m^2}\right)\frac{d\tilde{p}}{dh} \qquad (S10)$$

Equation (2b) allows us to divide by $\frac{\partial^2 w_P}{\partial p_m^2} + \frac{\partial^2 w_P}{\partial p_m \partial p}$ to get:

$$\left(\frac{\partial^2 w_H}{\partial h_m^2} + \frac{\partial^2 w_H}{\partial h_m \partial h}\right) < -\frac{\partial^2 w_H}{\partial h_m \partial p}\frac{d\tilde{p}}{dh} \qquad (S11)$$

which can simply be re-arranged to give equation (8):

$$\left(\frac{\partial^2 w_H}{\partial h_m^2} + \frac{\partial^2 w_H}{\partial h_m \partial h}\right) + \frac{\partial^2 w_H}{\partial h_m \partial p}\frac{d\tilde{p}}{dh} < 0 \qquad (8)$$

Equation (2b) is the same as equation (5), evaluated at the co-singular strategy. Therefore, if both conditions (2b) and (2c) hold, then both conditions (5) and (8) must hold.

Now instead assume that our co-singular strategy satisfies conditions (5) and (8) from the main text, at the co-singular strategy. That is:

$$\frac{\partial^2 w_P}{\partial p_m^2} + \frac{\partial^2 w_P}{\partial p_m \partial p} < 0 \tag{5}$$

$$\left(\frac{\partial^2 w_H}{\partial h_m^2} + \frac{\partial^2 w_H}{\partial h_m \partial h}\right) + \frac{\partial^2 w_H}{\partial h_m \partial p}\frac{d\tilde{p}}{dh} < 0 \tag{8}$$

Equation (5) allows us to multiply equation (8) by $\frac{\partial^2 w_P}{\partial p_m^2} + \frac{\partial^2 w_P}{\partial p_m \partial p}$ to get:

$$\left(\frac{\partial^2 w_H}{\partial h_m^2} + \frac{\partial^2 w_H}{\partial h_m \partial h}\right)\left(\frac{\partial^2 w_P}{\partial p_m^2} + \frac{\partial^2 w_P}{\partial p_m \partial p}\right) + \frac{\partial^2 w_H}{\partial h_m \partial p}\frac{d\tilde{p}}{dh}\left(\frac{\partial^2 w_P}{\partial p_m^2} + \frac{\partial^2 w_P}{\partial p_m \partial p}\right) < 0 \tag{S12}$$

Now we can use equation (S9) to re-write this as:

$$\left(\frac{\partial^2 w_H}{\partial h_m^2} + \frac{\partial^2 w_H}{\partial h_m \partial h}\right)\left(\frac{\partial^2 w_P}{\partial p_m^2} + \frac{\partial^2 w_P}{\partial p_m \partial p}\right) - \frac{\partial^2 w_H}{\partial h_m \partial p}\frac{\partial^2 w_P}{\partial p_m \partial h} < 0 \tag{S13}$$

This can simply be re-arranged to give equation (2c):

$$\left(\frac{\partial^2 w_H}{\partial h_m^2} + \frac{\partial^2 w_H}{\partial h_m \partial h}\right)\left(\frac{\partial^2 w_P}{\partial p_m^2} + \frac{\partial^2 w_P}{\partial p_m \partial p}\right) > \frac{\partial^2 w_H}{\partial h_m \partial p}\frac{\partial^2 w_P}{\partial p_m \partial h} \tag{2c}$$

Equation (5) evaluated at the co-singular strategy is the same as equation (2b). Therefore, if both conditions (5) and (8) hold, then both conditions (2b) and (2c) must hold. That is, conditions (2b) and (2c) combined are equivalent to conditions (5) and (8) combined. Either of these pairs of conditions can therefore be used to determine when a co-singular strategy is convergence stable, in the case where the parasite evolves much faster than the host.

#### RELATIONSHIP TO CONDITIONS GIVEN IN LEIMAR (2009)

Leimar (2009) states the following conditions for convergence to a co-singular strategy in a two-dimensional trait space:

$$\left(\frac{\partial^2 w_H}{\partial h_m^2}+\frac{\partial^2 w_H}{\partial h_m \partial h}\right)\left(\frac{\partial^2 w_P}{\partial p_m^2}+\frac{\partial^2 w_P}{\partial p_m \partial p}\right)-\frac{\partial^2 w_H}{\partial h_m \partial p}\frac{\partial^2 w_P}{\partial p_m \partial h}>0 \qquad (S14a)$$

$$A_{11}\left(\frac{\partial^2 w_H}{\partial h_m^2}+\frac{\partial^2 w_H}{\partial h_m \partial h}\right)+A_{22}\left(\frac{\partial^2 w_P}{\partial p_m^2}+\frac{\partial^2 w_P}{\partial p_m \partial p}\right)<0 \qquad (S14b)$$

where $A_{11}$ and $A_{22}$ are entries of the mutational matrix representing the mutate rates of the two host and parasite respectively [1]. Note that these are conditions (A1) and (A3) in the appendix of Leimar's paper. If we consider the case where the parasite mutates arbitrarily more quickly than the host (which implies that $A_{11} \ll A_{22}$), then these conditions reduce to:

$$\left(\frac{\partial^2 w_H}{\partial h_m^2}+\frac{\partial^2 w_H}{\partial h_m \partial h}\right)\left(\frac{\partial^2 w_P}{\partial p_m^2}+\frac{\partial^2 w_P}{\partial p_m \partial p}\right)-\frac{\partial^2 w_H}{\partial h_m \partial p}\frac{\partial^2 w_P}{\partial p_m \partial h}>0 \qquad (2c)$$

$$\frac{\partial^2 w_P}{\partial p_m^2}+\frac{\partial^2 w_P}{\partial p_m \partial p}<0 \qquad (2b)$$

which correspond to our conditions for convergence stability.

#### EXAMPLE 1 ANALYSIS

The endemic equilibrium of the system is given by:

$$S^*=\frac{\alpha+b}{\beta} \qquad (S15a)$$

$$I^*=\frac{\beta(a-b)-q(\alpha+b)}{\beta(\beta+q)} \qquad (S15b)$$

The invasion fitness of a mutant pathogen is given by:

$$w_P(p_m,p,h)=\frac{\beta(h,p_m)(\alpha+b)}{\beta(h,p)}-(\alpha+b) \qquad (S16a)$$

and the invasion fitness of a mutant host is given by:

$$w_H(h_m,p,h)=\frac{a(h_m)-qN^*(h,p)}{b+\beta(h_m,p)I^*(h,p)}-1 \qquad (S16b)$$

Let $\lambda=\frac{\beta_{0\max}-\beta_{0\min}}{p_0-p_1}$ and $\mu=\frac{a_1-a_0}{h_0-h_1}$ for notational convenience. Then the parasite singular strategy (when the parasite is evolving quickly relative to the host) is given by:

$$\hat{p}(h) = \frac{1}{2}p_1 + \frac{1}{2}h - \frac{\beta_{0\min}}{2\lambda} - 2 \qquad (S17)$$

and hence its derivative is:

$$\frac{\mathrm{d}\hat{p}}{\mathrm{d}h} = \frac{1}{2} \qquad (S18)$$

We also find that:

$$\left.\frac{\partial^2 w_P}{\partial p_m^2}\right|_{p_m=p=\hat{p}} = \frac{\lambda(\alpha + b)}{2\beta(p,h)} < 0 \qquad (S19)$$

This inequality holds for all values of $h$ and so the parasite is always evolutionarily stable, whether it is evolving quickly (in which case the above expression is in terms of $h$) or whether it is evolving on the same timescale as the host (in which case the above expression is evaluated at $h = h^*$). We also have that:

$$\left.\frac{\partial^2 w_P}{\partial p_m \partial p}\right|_{p_m=p=\hat{p}} = 0 \qquad (S20)$$

which implies that:

$$\left.\frac{\partial^2 w_P}{\partial p_m^2}\right|_{p_m=p=\hat{p}} + \left.\frac{\partial^2 w_P}{\partial p_m \partial p}\right|_{p_m=p=\hat{p}} = \frac{\lambda(\alpha + b)}{2\beta(p,h)} < 0 \qquad (S21)$$

This holds for all values of $h$ and so the parasite is always convergence stable when evolving quickly relative to the host.

We can also find the second derivatives of the host invasion fitness:

$$\left.\frac{\partial^2 w_H}{\partial h_m^2}\right|_{\substack{h_m=h=h^* \\ p=p^*}} = \frac{2\varepsilon}{b + \beta I^*} \qquad (S22)$$

This can be positive or negative, depending on the sign of the parameter $\varepsilon$ (which represents the curvature of the host trade-off), and so the host may be either evolutionarily stable or unstable (whether evolving alongside a fast pathogen or not). We also have:

$$\left.\frac{\partial^2 w_H}{\partial h_m \partial h}\right|_{\substack{h_m=h=h^* \\ p=p^*}} = \frac{q\beta_0^2}{16(\beta+q)(b+\beta I^*)^2}\left({I^*}^2 - I^*\frac{a+\alpha}{\beta+q} - \frac{b(\alpha+b)}{\beta^2}\right) \qquad (S23)$$

This means that $\left.\frac{\partial^2 w_H}{\partial h_m^2}\right|_{\substack{h_m=h=h^* \\ p=p^*}} + \left.\frac{\partial^2 w_H}{\partial h_m \partial h}\right|_{\substack{h_m=h=h^* \\ p=p^*}}$ can be positive or negative and so the convergence stability of the host (when the pathogen is evolving quickly) depends on the parameter values.

We can also calculate the rest of the second derivatives:

$$\left.\frac{\partial^2 w_P}{\partial p_m \partial h}\right|_{\substack{p_m=p=p^* \\ h=h^*}} = -\frac{(\alpha+b)\lambda}{4\beta} > 0 \qquad (S24)$$

$$\left.\frac{\partial^2 w_H}{\partial h_m \partial p}\right|_{\substack{h_m=h=h^* \\ p=p^*}} = \frac{\lambda I^*}{4(b+\beta I^*)} < 0 \qquad (S25)$$

This means that:

$$\left.\frac{\partial^2 w_H}{\partial h_m \partial p}\right|_{\substack{h_m=h=h^* \\ p=p^*}} \times \frac{\mathrm{d}\hat{p}}{\mathrm{d}h} = \frac{\lambda I^*}{8(b+\beta I^*)} < 0 \qquad (S26)$$

Therefore, strong convergence stability implies host fast convergence stability.

#### DESCRIPTION OF EVOLUTIONARY SIMULATIONS

1. Run the ecological dynamics of the system for a fixed length of time, with the host and parasite traits taking their resident values.
2. Introduce a mutant, randomly determining whether the mutation will occur in the host or parasite trait (the host mutates with probability $\frac{1}{1+\phi}$) and whether the trait will mutate to be slightly higher or lower than its current value. Add a small sub-population with the new, mutant trait values.
3. Run the ecological dynamics of the system for a fixed length of time, starting at its current composition.
4. Remove any sub-populations which have a density below a low threshold (extinct).
5. Introduce a mutant by randomly determining whether the mutation will occur in the host or parasite, whether the mutation arises in a susceptible host (and what trait value that host will take) or infected host (and what combination of host and parasite traits are present) and whether the mutating trait will mutate to be slightly higher or lower than its current value. Add a small sub-population with the new, mutant trait values.
6. Repeat steps 3 to 5 for many evolutionary timesteps.

| Parameter/ variable | Description | Default value or range |
|---|---|---|
| $\beta$ | Infectivity of pathogen to host | n/a |
| $\beta_0$ | Pathogen transmissibility | n/a |
| $\beta_{0\min}$ | Minimum value of pathogen transmissibility | 0 |
| $\beta_{0\max}$ | Maximum value of pathogen transmissibility | 15 |
| $p_0$ | Minimum value of pathogen evolving trait | 0 |
| $p_1$ | Maximum value of pathogen evolving trait | 1 |
| $h_0$ | Minimum value of host evolving trait | 0 |
| $h_1$ | Maximum value of host evolving trait | 4 |
| $a$ | Host reproduction rate | n/a |
| $a_0$ | Minimum value of host reproduction rate | 0.82 |
| $a_1$ | Host trait/reproduction trade-off parameter | 5.5 |
| $\alpha$ | Mortality virulence | 1.5 |
| $b$ | Host natural death rate | 0.5 |
| $q$ | Strength of host density dependence | 0.01 |
| $\varepsilon$ | Curvature of reproduction trade-off | 0.3 |
| $h, p$ | Host/parasite evolving traits | n/a |
| $S, I$ | Density of susceptible/infected hosts | n/a |
| $N$ | Total host density | n/a |
| $\phi$ | Relative pathogen mutation rate | n/a |

Table S1: Variables and parameters used in Example 1.

| Parameter/ variable | Description | Default value or range |
|---|---|---|
| $A$ | Host intrinsic mortality rate | 1 |
| $B$ | Host density-dependent mortality | 1 |
| $c$ | Transmission rate of highly virulent pathogen | n/a |
| $a$ | Virulence/transmission trade-off parameter | 10 |
| $K$ | Virulence/transmission trade-off parameter | 0.325 |
| $\bar{\alpha}$ | Virulence/transmission trade-off parameter | 6 |
| $\sigma$ | Spread of virulence/transmission trade-off | 10 |
| $c_0$ | Maximum value of transmission rate | 400 |
| $b_0$ | Maximum value of host reproduction rate | 12 |
| $c_1^b$ | Strength of resistance/reproduction trade-off | 0.8 |
| $c_2^b$ | Shape of resistance/reproduction trade-off | 5 |
| $\alpha$ | Parasite virulence (disease-induced mortality) | n/a |
| $r$ | Host resistance | n/a |
| $\beta$ | Transmission rate | n/a |
| $b$ | Host reproduction rate | n/a |
| $S, I$ | Density of susceptible/infected hosts | n/a |
| $N$ | Total host density | n/a |
| $\phi$ | Relative pathogen mutation rate | n/a |

Table S2: Parameters and variables used in Example 2.
